# Automated, high-throughput in-situ hybridization of *Lytechinus pictus* embryos

**DOI:** 10.1101/2025.03.23.644641

**Authors:** Yoon Lee, Chloe Jenniches, Rachel Metry, Gloria Renaudin, Svenja Kling, Evan Tjeerdema, Elliot W Jackson, Amro Hamdoun

## Abstract

Despite the reach of *in situ* hybridization (ISH) in developmental biology, it has rarely been used at scale. The major limitation has been the throughput of the assay, which typically relies upon labor intensive manual steps. The goal of this study was to develop a fully automated hybridization chain reaction (HCR) pipeline capable of large-scale gene expression pattern profiling, with dramatically reduced cost and effort, in the sea urchin *Lytechinus pictus*. Our resulting pipeline, which we term high throughput (HT)-HCR, can process 192 gene probe sets on whole-mount embryos within 32 hours. The unique qualities of the sea urchin embryo enabled us to automate the entire HCR assay in a 96-well plate format, and utilize highly miniaturized reaction volumes, a general purpose robotic liquid handler, and automated confocal microscopy. From this approach we produced high quality localization data for 101 target genes across three developmental stages of *L. pictus*. The results reveal the localization of previously undescribed physiological genes, as well as canonical developmental transcription factors. HT-HCR represents a log order increase in the rate at which spatial transcriptomic data can be resolved in the sea urchin. This study paves the way for localization of understudied genes and for sophisticated perturbation analysis.

**Summary Statement:** We developed an automated high-throughput HCR pipeline to rapidly map expression of 101 genes in sea urchin embryos, enabling large-scale discovery of novel developmental gene expression patterns.

## Introduction

*In situ* hybridization (ISH) is a foundational method in developmental biology. In sea urchins, it has been the basis for understanding cell fate, identity, and function (Juliano et al., 2006; Juliano et al., 2010; Luo and Su, 2012), as well as for the description of some of the most detailed developmental gene regulatory networks of development (Angerer and Angerer, 1981; Davidson et al., 2002). These studies take advantage of the tight development synchrony of this animal model, which makes it uniquely well suited for the study of spatial and temporal control of gene expression during development.

Yet, despite this long history in the field (Angerer and Angerer, 1981; Davidson et al., 2002), ISH in sea urchins remains a tedious manual procedure, often requiring years of effort to characterize large gene sets (Valencia and Peter, 2024). This has made the application of the technique prohibitive for large scale screens of gene expression patterns, for instance, as required to generate developmental gene expression atlases or for perturbation analyses. The advent of stable genetics in the sea urchin *Lytechinus pictus* (Jackson et al., 2024; Vyas et al., 2022) has paved the way for on-demand generation of millions of mutant, transgenic or knock-in embryos, which are useful for diverse studies from drug screens to perturbation studies. Yet without dramatic improvement in the throughput of in-situ hybridization, it could be impossible to take full advantage of the unique biological features (high fecundity, small embryo size, optical transparency, and developmental synchronicity) of sea urchins. To this end we leveraged the characteristics of the in-situ method hybridization chain reaction (HCR), to develop a fully automated, high-throughput ISH pipeline.

HCR is based on the use of affordable synthetic short (45nt) DNA oligonucleotide probes that innately suppress background and enable single step multiplexing (Choi et al., 2014; Choi et al., 2018). These qualities facilitate automation and increased throughput in two ways. First, HCR probes are easily commercially synthesized, within a week, as pools of unmodified oligonucleotides containing a set of short probes specific to several (typically three to five) gene targets, using a synthesis process comparable to primer synthesis. This allows researchers to quickly and affordably obtain probes for numerous targets, a vastly more efficient approach than production of long RNA probes. Second, automation of both multiplexing and hybridization incubation periods is made possible by the mechanism of signal amplification. Specifically, multiplexed signal amplification is enabled by using sequence-specific reagents to drive discrete hybridization chain reactions of different channels of fluorescent oligonucleotides specific to their respective gene targets. This eliminates the need for many of the sequential multiplexing steps which are common to other ISH methods. In addition, the amplification of the signal is driven by sequence complementarity of fluorescent oligonucleotides in a controlled molecular crowdant buffer, reducing the possibility of nonspecific signal and overstaining.

The goal of this study was to develop a robust pipeline for high-throughput sea urchin spatial transcriptomics, that can be easily applied by end users. Prior efforts to automate in-situ hybridization have typically relied upon bespoke hardware solutions, with limited throughput and utility in other assays. Our approach relies on plate and robotic formats likely to be found in many core facilities or individual labs. These general liquid handlers require smaller capital expenditure risk, since the instrument can also perform a variety of other routine tasks. Furthermore, the capability to complete the assay in standard plate formats allow for the automation of image data acquisition using automated confocal imaging systems commonly used in high-content drug screening assays (Boutros et al., 2015; Dranchak et al., 2023; Stossi et al., 2023).

Here, we present high-throughput HCR (HT-HCR), a fully automated, miniaturized HCR pipeline capable of running 192 probe sets in 32 hours. Using HT-HCR we probed three early developmental stages of *L. pictus* embryos, using a probe panel designed to target genes expressed at different developmental stages, in different spatial domains, and representing a wide range of functional categories. The results resolved the localization patterns of 101 genes in *Lytechinus pictus* belonging to multiple KEGG categories (Kanehisa, 2019; Kanehisa and Goto, 2000; Kanehisa et al., 2023), including cell motility (2), cytoskeleton (4), genetic information processing (5), matrix proteins (3), membrane trafficking (3), membrane transport (18), metabolism (5), signal transduction (12), transcription factors (31), transport and catabolism (2), and 16 genes categorized as “other”.

## Methods

### Probe design

Probe sets for 192 targets predicted in the genome assembly UCSD_Lpic_2.1 were designed using a version of insitu_probe_generator (Kuehn et al., 2022) containing minor modifications to facilitate the design of probes in multiplexed oligonucleotide pools, automatic retrieval of target information and metadata from NCBI, input of target coding sequences (CDS), and generation of metadata which tracks and labels intermediate outputs of the probe pool design process. These were necessary for data management and image processing pipeline.

Individual probes were all designed to have a maximum homopolymer length of three nucleotides. The maximum number of probe pairs (i.e. probe set) for a given target gene was 15. Because the number of probe pairs depends on the length of the target gene, the number of probe pairs per target gene varied from 3 to 15. Each probe set was designed to be specific to one of four different amplifier sequences, B1, B2, B3, and B4 to multiplex samples in a probe pool for four gene targets (Choi et al. 2018). Thus, each probe pool contained four probe sets. A total of 48 probe pools were ordered and synthesized (Integrated DNA Technologies, Coralville, Iowa, USA) at a scale of 100 pmol per oligonucleotide sequence. Oligonucleotide pools were resuspended in 100 µ L of 1X TE buffer or nuclease-free water to obtain a 1 µM concentration per oligonucleotide per pool and were stored in -20°C. All probe sequences designed for this project are available in the Supplementary Materials.

### Embryo culturing and fixation

Three independent batches were run for 24 hpf embryo samples, and three batches were run for mixed stage samples. For each batch of embryos, sperm from one male and eggs from two to four female *L. pictus* adults were spawned and mixed according to (Nesbit and Hamdoun, 2020).

Eggs were washed with 0.2 µm filtered sea water (FSW) and any debris collected while spawning was removed. After fertilizing the eggs, the culture density was normalized to 500 embryos/mL in 2 L beakers. Cultures were raised at 21°C. Embryos were collected at 12, 24, and 36 hours post fertilization (hpf) by aspirating off the surface with a 50 mL serological pipette and concentrating the embryos on a 75 µm strainer. The embryos were then transferred to a 15 mL tube and allowed to settle on ice. Next, the supernatant was removed until 2 mL of embryos in FSW remained. This was transferred to a 2 mL cryogenic tube, and the concentration of embryos was approximately 250,000 embryos/mL. Concentrated embryos were placed on ice for 10 minutes to allow the embryos to settle to the bottom of 2 mL tubes. This step was repeated in between all steps that involve agitation or mixing of the embryos. Once settled, 1 mL of the supernatant was removed, and 1 mL ice-cold 1M NaCl was added to bring the sample to a final concentration to 0.5M NaCl to deciliate the embryos. Following deciliation, as much supernatant as possible was first removed, followed by two 1.5 mL washes with FSW. After the final FSW wash, 1 mL of the supernatant was removed, and 1 mL of 8% paraformaldehyde was added to 1 mL of the embryos, bringing the final paraformaldehyde concentration to 4%. Samples were incubated at 4°C on a rocker for 48 hours. One important deviation to note here from previously published HCR protocols is that ice, not centrifugation, was used to concentrate the samples for fixation preparation. The reason being is that centrifugation inevitably introduces undesirable debris and clumping into the final prepared sample.

After incubation in 4% paraformaldehyde, the samples were kept on ice and washed three times with 1 mL 1X phosphate buffered saline with a final concentration of Tween 20 at 10%, with 10 minute incubation time in between all washes. Subsequently, three washes were performed using 1 mL of 50%, 75%, and 100% methanol accordingly. After the final methanol wash, each 2 mL sample was topped off to 2 mL with 100% methanol, bringing each sample tube to approximately 40-60 embryos/µL. Samples were stored in -20°C for use in the automated HCR assay.

### Consumable and reagent preparation

All consumable plates and reagents were prepared according to the final concentrations found in the HCR RNA-FISH protocol for whole-mount sea urchin embryos (Molecular Instruments Incorporated, Los Angeles, California, USA). All bulk reagents, probe wash buffer, amplification buffer, and 5X saline sodium citrate (SSCT) with a final concentration of Tween 20 at 10% were aliquoted into 12-well, 15 mL trough plates. These bulk reagent plates were heat sealed and stored at 4°C. The first well of each 12-well trough plate was left empty to be loaded with rehydrated fixed embryos.

The following describes the sample rehydration process performed immediately prior to starting all runs. Fixed embryos in methanol were left upright on ice to settle to the bottom of the 2 mL tube. 1.7 m L of methanol were removed from the sample. Five 1.5 mL washes are performed using 5X SSCT at room temperature. For each wash, samples were incubated until embryos settled to the bottom of the tube. After the final wash, leaving a total of 300 µL of embryos in 5X SSCT, 800 µL of warmed (37°C) hybridization buffer is added to the sample. The sample is left to pre-hybridize at 37°C for at least one hour. Pre-hybridized embryos were transferred to the empty first well of the bulk reagent plate immediately before the run.

Primary stock probe plates were created by mixing 10 µL from a 1 µM stock probe pool solution with 90 µ L of hybridization buffer (HB) into each well of a 96-well plate according to a predefined plate map, bringing the probe pool concentration in each well to 100 nM. An intermediate probe plate was then created by mixing 20 µL from the primary stock probe plate and 80 µL of hybridization buffer to obtain 100 µL of a 20 nM probe solution in each well. The contents of the intermediate probe plate were distributed across 20 consumable probe plates in 5 µL aliquots. This consumable plate is then used directly in the automated assay and discarded after use. All data in this publication have been produced using the same batch of probe plates that have been stored in this way for at least three months.

The amplifier hairpins used were B1H1/H2-Cyanine7, B2H1/2-Alexa Fluor 647, and B3H1/H2-Alexa Fluor 594 (Molecular Instruments Incorporated, Los Angeles, California, USA). We synthesized B4H1/H2 amplifier hairpins by conjugating amine-modified oligonucleotides with the B4 sequences with NHS-ester modified ATTO 532 and subsequently purifying the conjugated product using 15% urea-polyacrylamide gel purification (amine-modified oligonucleotide synthesis: Integrated DNA Technologies, Coralville, Iowa, USA; fluorophore: ATTO-TEC GmbH, Siegen, Germany). The resulting B4H1/2-ATTO 532 amplifier hairpins were normalized to 3µM. To make a stock hairpin master mix, 100µL of each amplifier hairpin at 3µM were incubated at 95°C for 90 seconds and snap-cooled at room temperature for one hour minimum. After incubation, all amplifiers are mixed into a single tube. The concentration of each hairpin at this stage is 375 nM. 25µL of this master mix was then distributed into each well of the first column across four 96-well plates to be used as consumable amplifier plates. All consumable reagent plates were heat sealed and stored at 4°C.

### Assay automation and miniaturization

Automation and miniaturization of HCR resulting in HT-HCR was accomplished using the Opentrons Flex (Opentrons Labworks Inc., Long Island City, New York, USA), a general-purpose laboratory liquid handler. HT-HCR is a miniaturized adaptation of the standard HCR protocol using reaction volumes reduced from 100µL to 15µL. The file encoding this protocol, *HL_HTHCR*, including the offsets from calibration for the modules and consumables, is available at https://github.com/ylee-sio/HL_HTHCR. The modules installed for this protocol included two Flex 8-Channel Pipettes (1-50 and 5-1000 µL), Flex Gripper (robotic arm), Opentrons Thermocycler GEN2, Heater-Shaker Module (with the Universal Flat Adapter), Temperature Module (with the PCR Plate Adapter), and the Flex Waste Chute. Robotic calibration was performed for each module and for the dimensions of the following consumables: full skirted 96-well PCR Plate (ThermoFisher Scientific, AB-3396), 96-deep well plate (Agilent, 5043-9300), 96-well glass-bottom imaging plate (Cellvis, P96-1.5H-N), and 12-well/trough 15mL reservoir (NEST, 360102).

A consumable probe plate was removed from -20°C, centrifuged, unsealed, and loaded onto the thermocycler module set at 37°C. The consumable amplifier plate was removed from 4°C, centrifuged, unsealed, and loaded onto the temperature module at 12°C. The temperature module was set to be 1-2°C above the temperature of the local dew point to prevent condensation on the bottom of the module. 1 mL of prehybridized embryos was dispensed into the empty trough left for loading fixed samples. The bulk reagent plate was then loaded onto the deck at room temperature.

Using the 8-channel pipette module, 10 µL of embryos were transferred from the bulk reagent plate to the consumable probe plate on the thermocycler and mixed vigorously. Following this, the thermocycler was set to close and seal the plate with the lid set at 40°C. This temperature was set to avoid condensation on the thermocycler seal. The embryos in the consumable probe plate were left to hybridize for 12 hours. After hybridization, samples were washed three times using 135 µL of probe wash buffer at 37°C, with a 30 minute settling period following each wash. Following this, samples were washed three times using 135 µL of 5X SSCT at 21°C, with a 30 minute settling period following each wash. After the final 5X SSCT, the plate contained 15 µL 5X SSCT with embryos settled at the bottom of each well. Samples were resuspended using 30 µ L of amplification buffer. Before amplification, 35 µL of samples suspended in amplification buffer were moved to the unused columns of the consumable amplifier plate, bringing the volume of samples in the probe plate down to 10 µL.

70 µL of amplification buffer was added to each well of the 25 µL master mix resulting in a concentration of 98 nM per probe. 12 µL of this diluted hairpin solution was transferred to the samples in the consumable probe plate, resulting in a final concentration of 53 nM per hairpin. Samples were incubated in hairpin solution for 12 hours at 21°C. After amplification, samples were washed using 135 µL 5X SSCT at 21°C, leaving 30 minutes between each wash for embryos to settle. Samples were then incubated in 135 µL 5X SSCT containing Hoechst 33342 for nuclear staining at 10 µg/mL at 21°C for 30 minutes. This nuclear stain in 5X SSCT was then removed, and a 135 µL 5X SSCT was added to resuspend the embryos. Immediately after resuspension, 135 µ L of the solution containing the resuspended embryos were transferred to a glass-bottom imaging plate sitting on a shaker module. After completing all transfers to the imaging plate, embryos were centered in their wells by shaking the plate at 500 rpm for 5 minutes to concentrate embryos in the center of the well and prepare them for imaging.

### Imaging

The imaging plate was loaded into the ImageXpress HT.ai Confocal Microscope (IXM HT.ai; Molecular Devices, San Jose, California, USA). Samples were imaged with a 20X 0.8NA Plan Apo Lambda objective (Nikon, Minato City, Tokyo, Japan) with the following configurations for the laser light source (89North, Williston, Vermont, USA) and emission filters (Semrock Incorporated, Rochester, New York, USA) combinations: 740nm with a 794/32 filter for Cyanine7, 640nm with a 680/42 filter for Alexa Fluor 647, 555 nm with a 624/40 filter for Alexa Fluor 594, 520nm with a 562/40 filter for ATTO 532, and 405nm with a 452/45 filter for Hoechst 33342. Exposure times were set for 150ms for all channels except for the 405nm channel, which was set at 50 ms due to the higher intensity of the nuclear stain signal.

For a standard circular bottom image plate, there were a total of 81 sites (9 by 9 grid) available to image using a 20X objective. However, only the center 4-9 sites (2x2 or 3x3) were imaged for data storage efficiency. For images with individual stages of embryos, z-slices were taken in 2 µm intervals. 35 slices were taken for 12 hpf samples, 40 slices were taken for 24 hpf samples, and 55 slices were taken for 36 hpf samples. This covers 70 µm, 80 µm, and 110 µm for 12, 24, and 36 hpf samples, respectively. The total duration for imaging 48 wells of samples ranged between 3-4 hours depending on the developmental stage of the sample. A total of 75,600 (151.2GB), 86,400 (172.8GB), and 118,800 (237.6GB) images were generated for 12 hpf, 24 hpf, and 36 hpf runs, respectively (# sites x # slices x # wells x # channels), for nine (3x3) sites.

In order to reduce assay run times, reagents, consumables, data storage, and image processing time, we opted for performing the assay for developmental stage comparisons using mixed stage samples. For assay runs containing mixed stage samples, equal parts of fixed embryos from the three different developmental stages were mixed at the rehydration step. No other changes were made for running the assay for mixed stage samples. For images containing mixed developmental stages, 45 z-slices were taken in 2 µm intervals, resulting in 97,200 images (194.4GB). All imaging was performed at room temperature (20-23°C).

### Data management and processing

Custom scripts for automatic offloading and archival management of data from the IXM HT.ai acquisition software (MetaXpress 6 Software) were written in bash, Microsoft Powershell, and Python. Scripts for automated metadata annotation and application (target text annotation on images, pseudocolor assignment, channel multiplexing and demultiplexing, aspect ratio adjustment according hardware binning levels, site tiling, and directory management tasks) were written in bash, R, and ImageJ Macros (Schneider et al., 2012). These scripts are available at https://github.com/ylee-sio/HL_HTHCR.

After data acquisition of all samples, images were automatically sorted by well ID (which links to an oligo pool ID), site ID, and wavelength (which links to the hairpin amplifier sequence and fluorophore used for the localization of a specific gene). The 45 images of each z-slice from each site for each wavelength were maximally projected and the center site image, which typically contained the greatest number of embryos within a site among all sites, was used to determine whether the probe produced a positive signal. Generally, we selected images with signal that was visually discernible from background (signal from non-sample area). For presentation of images from 101 probes with positive localization results, images were either stitched to form either the full 3x3 or 2x2 tiled images or used as single sites. Information linked to well IDs and wavelengths were used to automatically annotate each image with a common gene name and NCBI accession for the specific mRNA sequence used to design the probe set.

## Results

Our HT-HCR screen produced localization data for 101 genes in *Lytechinus pictus*. The images are publicly available on our laboratory data repository which is available through the following link <u>Hamdoun Lab Image</u> <u>Repository</u>. These included genes with a broad range of biological functions and localization patterns along the KEGG BRITE in *L. pictus* (**Figure S1, Table S1**).

### Validation of the automated assay

To test the robustness of our automated HCR protocol (**Figure 1**), we optimized our liquid handling steps for accurate miniaturized sample volumes and maintenance of sample morphological integrity. Our optimization also considered efficient consumable usage (i.e. tip reuse steps) without the risk of sample contamination events. This was important for a fully automated protocol on a basic liquid handler because deck space limitations typically require users to be present to refill consumables during a paused state during the protocol run. We compared the results of the finalized protocol to those produced by manual HCR across stages and genes. One of our concerns was the maintenance of morphological integrity using automated liquid handling. We found that unmodified automation tips were able to transfer embryos in dense concentrations in a viscous hybridization buffer with little to no damage to 24 hpf embryos, a stage which is commonly used to study gastrulation (**Figure 2**). Furthermore, we tested the automated HCR protocol across different developmental stages (**Figure S2**) using probes for the genes *white protein* (**Figure S2A**), *collagen 1-alpha(V) chain* (**Figure S2B**), and *tubulin alpha 1 (tuba1)* (**Figure S2C**).

**Figure 1.**
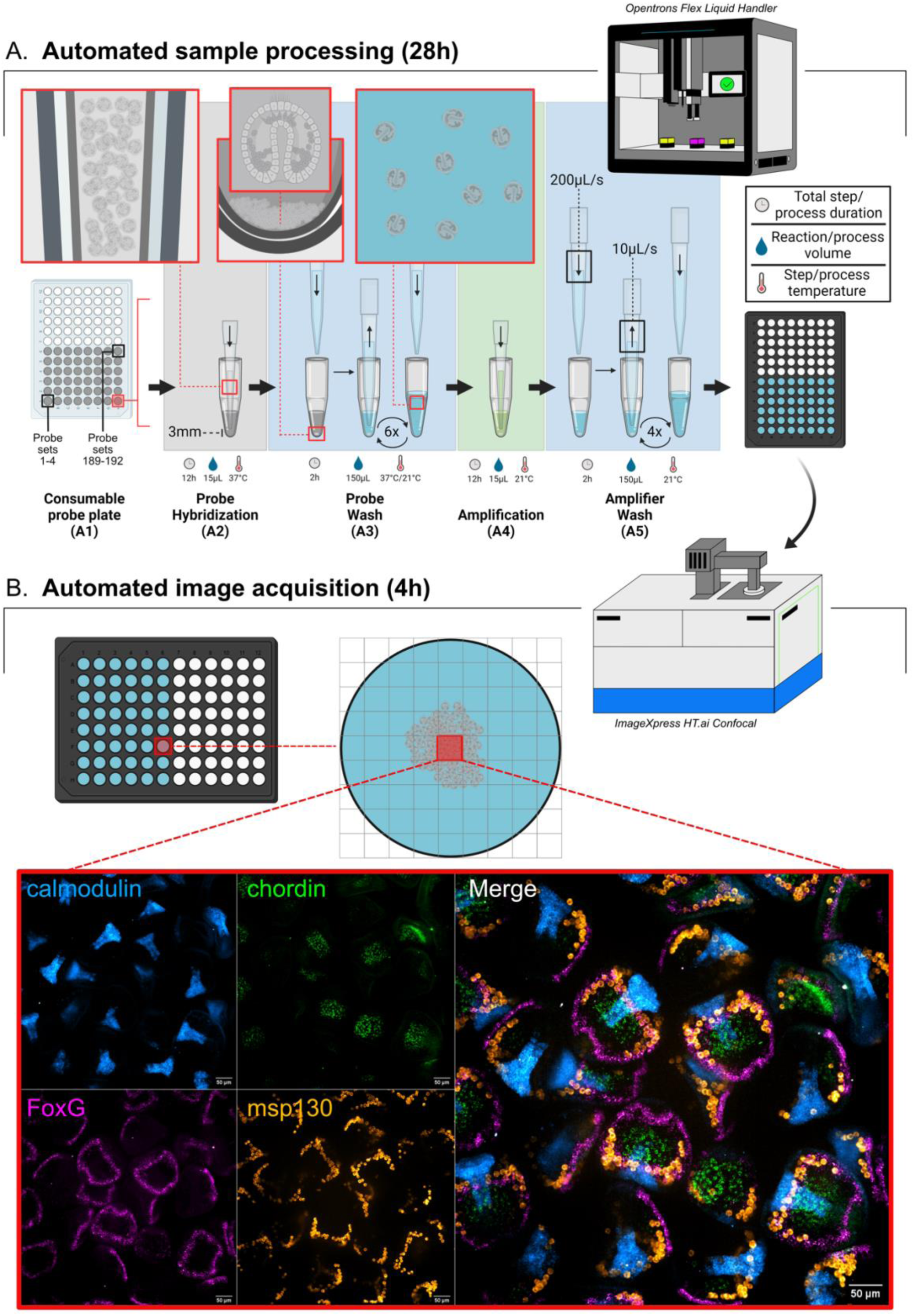
Diagram of a fully automated HCR workflow. A) Automated sample processing using the Opentrons Flex robot. A1) *Consumable probe plate as the starting vessel.* A 96-well plate, where each well contains 5μL of a pool of probes for five mRNA targets is used as the starting vessel for sample processing. For all liquid handling steps throughout the protocol, dispense speeds are set at 200 μL/s and aspiration speeds at 10 μL/s. All aspirations are set to use a tip to well bottom distance of 3mm. A2) *Probe hybridization step.* 10μL of fixed embryos in hybridization buffer are transferred to each well to start the probe hybridization step. The inset illustrates transferability of sea urchin embryos using standard automation pipette tips. A3) *Probe wash step.* Probes are washed from the sample using a formamide based probe wash buffer. The insets illustrate the liquid handling speeds and tip to bottom distances allow for resuspension and mixing of embryos throughout washes. A4) *Amplification step.* 10 μL of embryos are transferred to a plate of pre-plated amplifier hairpins in dextran sulfate. A5) *Amplifier wash step*. Amplifier hairpins are washed using 5X SSCT, and 150 μL of samples in 5X SSCT are subsequently transferred to a glass bottom high-content imaging plate. Samples are centered using a plate shaker module set at 450 rpm for 30 minutes. B) *Automated confocal image acquisition using the ImageXpress HT.ai.* The imaging plate containing centered samples are placed into the confocal microscope. Depending on data storage availability, the center 4 or 9 are imaged for each well. An example of an output of the demultiplexed and multiplexed versions of a single site of a positive control sample well is shown.

**Figure 2.**
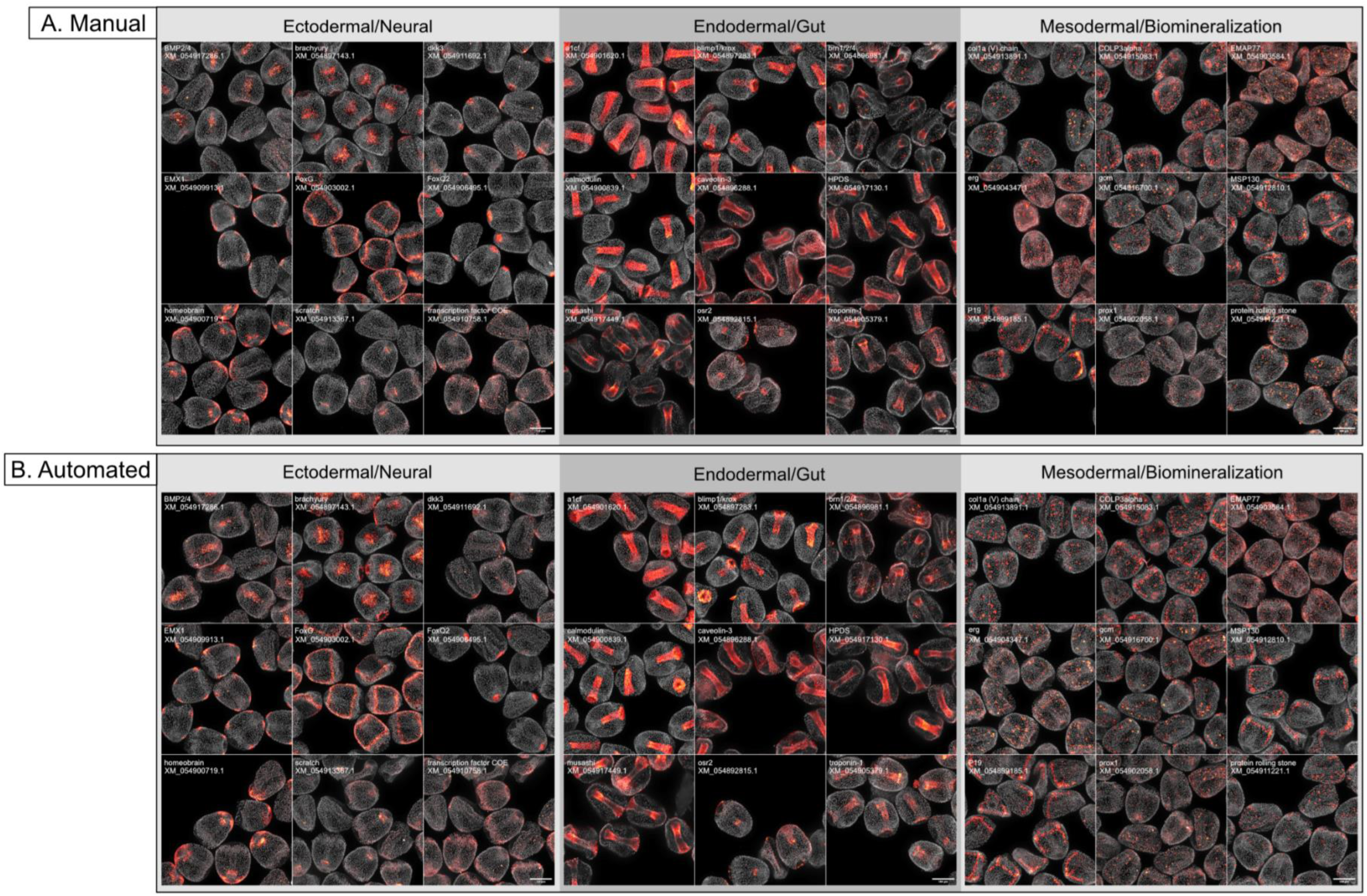
Comparison of results from manual and automated assays. Images show the localization of several positive control targets in 24 hpf embryos. Manually processed samples and robotically processed samples appear identical. A) Imaging results of manually processed samples. B) Imaging results of robotically processed samples. Nuclear staining using Hoescht 33342 appears in gray.

To determine whether our automated assay would be able to produce similar results to manual approaches across spatial domains, we compared outputs of our assay to those for 27 genes (**Figure 2**). We found that, qualitatively, the automated assay produced results comparable to those obtained by a manual approach. A comparison of the localization of three genes *empty spiracles homeobox 1* (*EMX1*), *hematopoietic prostaglandin D-synthase* (*HPDS*), and *P19* (previously shown by (Costa et al., 2012) across three different automated runs indicated that these results were produced consistently across batches and runs (**Figure S3**).

Next, we examined the efficacy of our automated protocol for mixed stage samples, focusing on transcription factors which were well-described in urchins and expressed in highly spatially restricted patterns (**Figure 3**). Overall, we found that unique morphological features of these three developmental stages allowed for easy differentiation of embryos across stages. Blastula stage embryos (12 hpf) were hollow spheres of cells, gastrula stage embryos (24 hpf) contained gut tubes, and prism stage embryos (36 hpf) possessed two distinct arms. Given this, we proceeded to perform the automated assay using mixed stage samples to conserve reagents, consumables, and number of runs.

**Figure 3.**
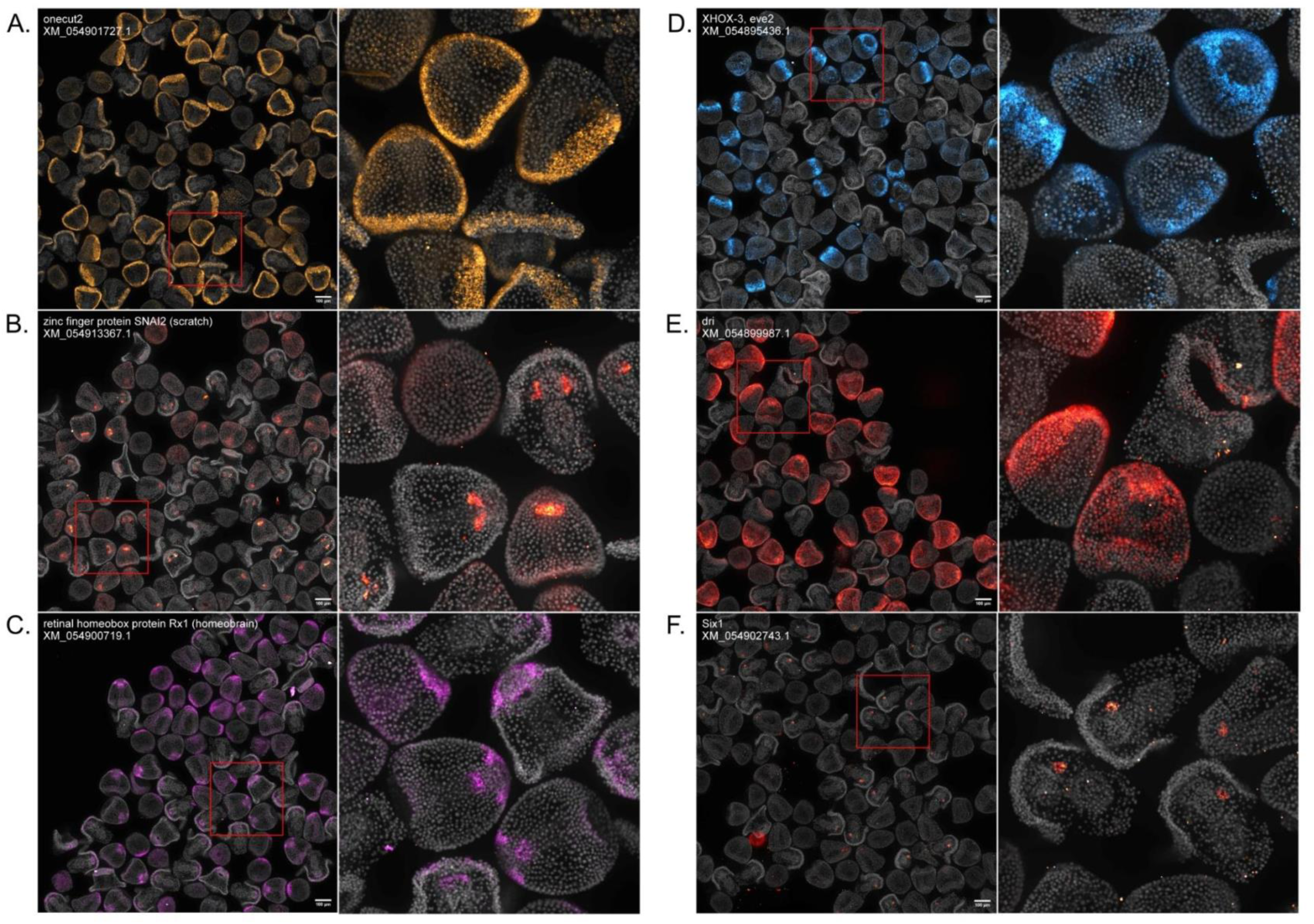
Validation of HT-HCR by localization of previously described transcription factor genes. All images contain a mix of 12, 24, and 36 hpf embryos and are automatically demultiplexed and stitched outputs resulting from an automated image processing workflow. Each image contains between 50-100 embryos and are annotated with the NCBI RefSeq ID for the mRNA sequence targeted by probe oligonucleotides. A) *onecut2:* Expression is sparse in 12 hpf embryos. In 24 hpf and 36 hpf embryos, expression is concentrated in the ciliary band. B) *scratch*: Expression is either sparse or absent in 12 hpf embryos. At 24 hpf, expression is concentrated in the archenteron. At 36 hpf, expression is restricted to the coelomic pouches. C) *homeobrain:* At 12 hpf, expression is present in the apical ectoderm. At 24 and 36 hpf, expression is in the apical ectoderm and stomodeum. In more developed 36 hpf, expression can be seen in the upper region of the developing oral hood. D) *XHOX-3, eve2:* Expression is most enriched in 12 hpf embryos in a ring of cells in the aboral ectoderm. Sparse expression is visible in some 24 hpf embryos. E) *dri:* Expression is most enriched in 24 hpf embryos and is restricted to the oral ectoderm. Sparse expression is visible along the ciliary band in 36 hpf embryos. F) *Six1:* Expression is restricted to the coelomic pouches in 36 hpf embryos.

Three of the localized transcription factors included *onecut2*, *scratch,* and *homeobrain* (**Figure 3A-C**) which are associated with neurodevelopment. The expression of *onecut2* is concentrated in the ciliary band in 24 and 36 hpf embryos but is sparse in 12 hpf embryos (**Figure 3A**). These patterns are similar to the localization patterns found in *Strongylocentrotus purpuratus* (Poustka et al., 2004). The expression of *scratch* in *L. pictus* appears different from the localization pattern found in *Lytechinus variegatus (Slota et al., 2019)* which showed expression that was concentrated in a few apical cells during at 24 hpf. In *L. pictus*, the expression of *scratch* appears to be absent in 12 hpf embryos and present in the foregut at 24 hpf and the coelomic pouches at 36 hpf (**Figure 3B**). This could be due to some species-specific difference in localization or developmental timing. The expression of *homeobrain* recapitulates the localization patterns found in another urchin species *Hemicentrotus pulcherrimus* by Yaguchi et al. 2016 (Yaguchi et al., 2016). *Homeobrain* is concentrated in the apical ectoderm in 12 hpf embryos (**Figure 3C**). At 24 hpf and 36 hpf, expression is in the apical ectoderm and stomodeum. In more developed 36 hpf, expression can be seen in the upper region of the developing oral hood.

Next, we localized several transcription factors whose expression patterns were enriched in highly specific timepoints. These included the expression patterns of *XHOX-3/eve2* (Peter and Davidson, 2010), *Deadringer (dri) (Amore et al., 2003)*, and *Six1 (Byrne et al., 2018)* (**Figure 3D-F**). *XHOX-3/eve2* expression is restricted to a ring of cells in the aboral ectoderm in 12 hpf embryos (**Figure 3D**). In 24 and 36 hpf embryos, expression is faint or absent. *Deadringer (dri)* expression is highly concentrated in the oral ectoderm in 24 hpf embryos (**Figure 3E**). Expression is absent in 12 hpf embryos, and in 36 hpf embryos, some faint expression is visible in the ciliary band. The expression of *Six1* is clearly visible in 36 hpf embryos in one of the coelomic pouches, and in 12 and 24 hpf embryos, the expression pattern is either sparse or absent (**Figure 3F**). Overall, these results indicate that HT-HCR is robust across spatiotemporal domains.

### Expression pattern discovery using the automated assay

An application of automated HCR is localization of understudied genes, at reduced cost and effort. Our screen revealed the expression patterns of several genes which, to our knowledge, have not been previously localized in urchins (**Figure 4**). These include genes important for cellular structure such as *gelsolin-2* and *mucin-2* (**Figure 4A**), enzymes such as *D-dopachrome decarboxylase* and *hematopoietic prostaglandin D-synthase* (**Figure 4B**), as well as new transcription factors such as *thyroid transcription factor-1* and *odd-skipped related-1* (**Figure 4C**). We also applied this tool to localization categories of genes that are challenging to localize by *in-situ* hybridization due to the presence of multiple related homologs and/or relatively low expression levels. Examples were ion channels and transporters, including ATP Binding Cassette (ABC) transporters and solute carriers (SLCs), as well as other membrane transporters belonging to additional gene families.

**Figure 4.**
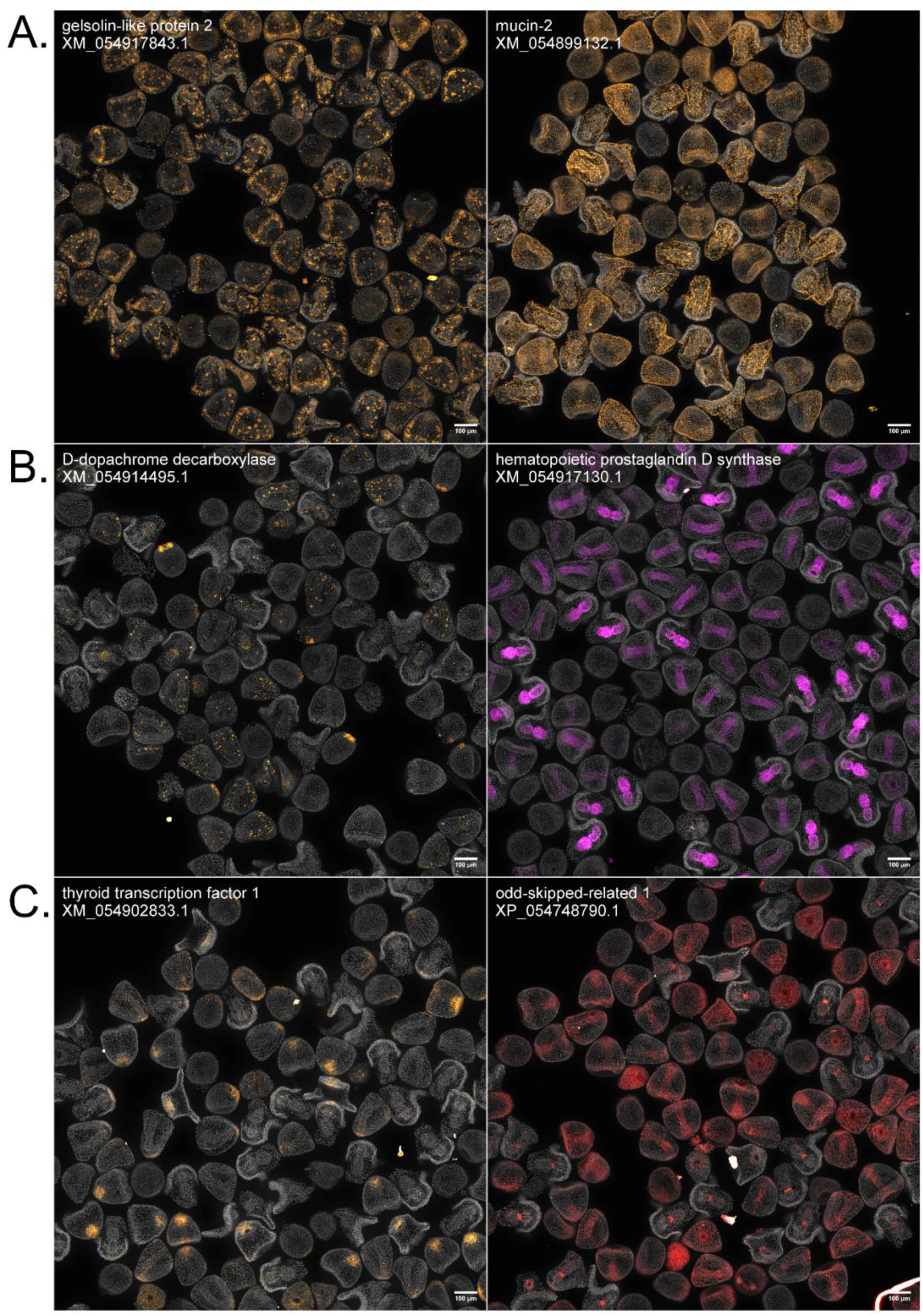
HT-HCR to localize previously undescribed cellular structure, enzyme, and transcription factor genes. All images contain a mix of 12, 24, and 36 hpf embryos and are automatically demultiplexed and stitched outputs resulting from an automated image processing workflow. Each image contains between 50-100 embryos and are annotated with the NCBI RefSeq ID for the mRNA sequence targeted by probe oligonucleotides. A) Genes involved in cellular structure: *gelsolin-like protein 2* and *mucin-2*. *Gelsolin-like protein 2*: Expression does not appear to be present in 12 hpf embryos. In 24 hpf embryos, expression is visible in PMCs and SMCs. *Mucin-2*: Expression is faint or absent in 12 hpf embryos. Expression is visible in the ectoderm in 24 and 36 hpf embryos. B) Enzymes: *D-dopachrome decarboxylase* and *hematopoietic prostaglandin D synthase*. *D-dopachrome decarboxylase*: Expression is visible in the NSM in 12 hpf embryos. Expression is visible in mesodermal cells in 24 hpf embryos. Expression is faint in 36 hpf embryos-some samples show signal in mesodermal cells and the midgut. *Hemtaopoietic prostaglandin D-synthase*: Expression is not present in 12 hpf embryos. In 24 hpf embryos, expression is visible in the gut tube. By 36 hpf, expression is localized in the midgut and foregut. C) Transcription factors: *thyroid transcription factor 1* and *odd-skipped related 1 protein. Thyroid transcription factor 1*: Expression is either faint or absent in 12 hpf embryos. In 24 hpf embryos, expression is visible in the apical ectoderm. In 36 hpf embryos, expression is visible in the oral hood. *Odd-skipped related 1 protein*: Expression is faint or absent in 12 hpf embryos. In 24 hpf embryos, expression appears to concentrate in the gut tissue. By 36 hpf, expression is restricted to the cardiac sphincter.

For ABC transporters (**Figure 5A-C**), new localizations included heme transporter *ABCB7* (Allikmets et al., 1999; Pondarré et al., 2006; Pondarre et al., 2007; Taketani et al., 2003) and lipid transporter *ABCA3* (Ban et al., 2007; Klugbauer and Hofmann, 1996; Schmitz and Langmann, 2001; Xie et al., 2022). The localization of lipid/sterol transporter *ABCG12* (Chen et al., 2011; McFarlane et al., 2010; Pighin et al., 2004) has previously been demonstrated in *S. purpuratus* (Lee et al., 2023) and was repeated for *L. pictus* in this study. *ABCB7* expression is ubiquitous in 12 hpf embryos. By 24 hpf, expression is still ubiquitous but more concentrated in what appear to be mesodermal cells (based on their location within the embryo), and by 36 hpf, *ABCB7* becomes more consistent with mesodermal cell expression (**Figure 5A**). *ABCA3* expression is ubiquitous in both 12 hpf and 24 hpf embryos, but by 36 hfp, the expression is restricted to the midgut (**Figure 5B**). The expression pattern for *ABCG12* followed the stereotypical spatiotemporal expression pattern of mesodermal genes (**Figure 5C**). 12 hpf embryos showed concentrated expression in the non-skeletogenic mesoderm (NSM) ring and by 24 hpf and 36 hpf, expression was concentrated in mesodermal cells.

**Figure 5.**
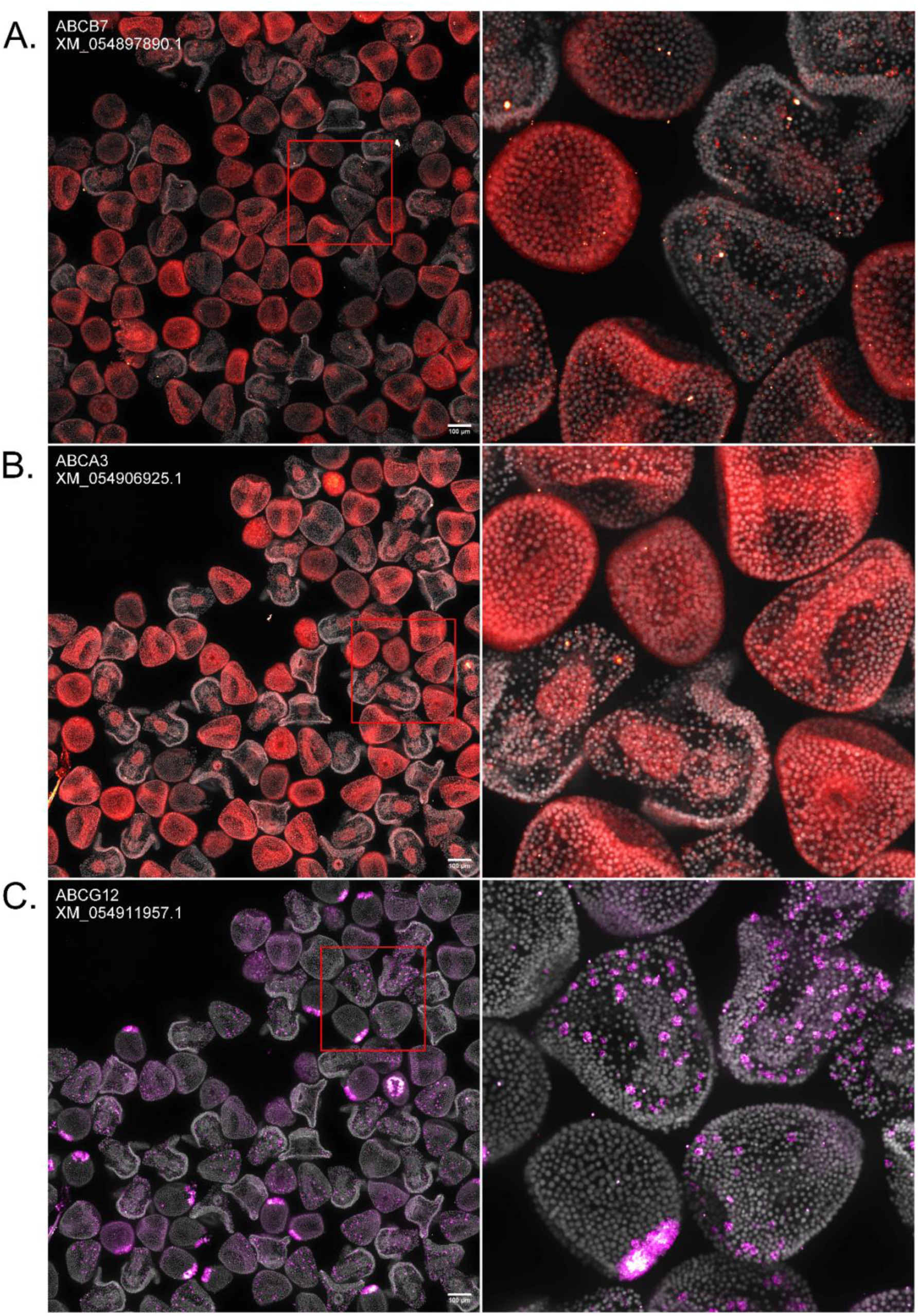
Novel ABC transporter expression patterns revealed by HT-HCR. All images contain a mix of 12, 24, and 36 hfp embryos and are automatically demultiplexed and stitched outputs resulting from an automated image processing workflow. Each image contains between 50-100 embryos and are annotated with the NCBI RefSeq ID for the mRNA sequence targeted by probe oligonucleotides. A) *ABCB7:* Expression is low in 12 hfp embryos. Expression is somewhat ubiquitous but concentrated in mesodermal cells in 24 hfp embryos. Expression is restricted to pigment cells by 36 hfp. B) *ABCA3:* Expression is ubiquitous in 12 hfp and 24 hfp embryos. Expression is concentrated in the midgut in 36 hfp embryos. C) *ABCG12:* Expression is concentrated in the NSM in 12 hfp embryos. At 24 hfp and 36 hfp, expression is concentrated in mesodermal cells.

For SLCs, new localizations included choline transporter *SLC5A7 (Apparsundaram et al., 2000; Iwamoto et al., 2006; Okuda and Haga, 2000)*, vesicular acetylcholine transporter SLC18A3 (Varoqui and Erickson, 1996; Weihe et al., 1996), neutral and dibasic amino acid transporter *SLC6A14* (Anderson et al., 2008; Sloan and Mager, 1999) (**Figure 6A-C**). At 12 hpf, the expression of *SLC5A7* is sparse or absent (**Figure 6A**). In 24 hpf embryos, expression is restricted to cells which reside in the putative location of post-oral neural progenitor cells described in (Slota et al., 2019). In 36 hpf embryos, expression was sparse or absent (**Figure 6A**). The expression pattern of *SLC18A3* also resembles those of genes expressed in post-oral neural progenitor cells at 24 hpf (**Figure 6B**). By 36 hpf, *SLC18A3*^+^ cells are scattered along the ciliary band. *SLC6A14* expression is sparse or absent in 12 hpf and 24 hpf, but by 36 hpf, the expression is restricted to one of the coelomic pouches (**Figure 6C**).

**Figure 6.**
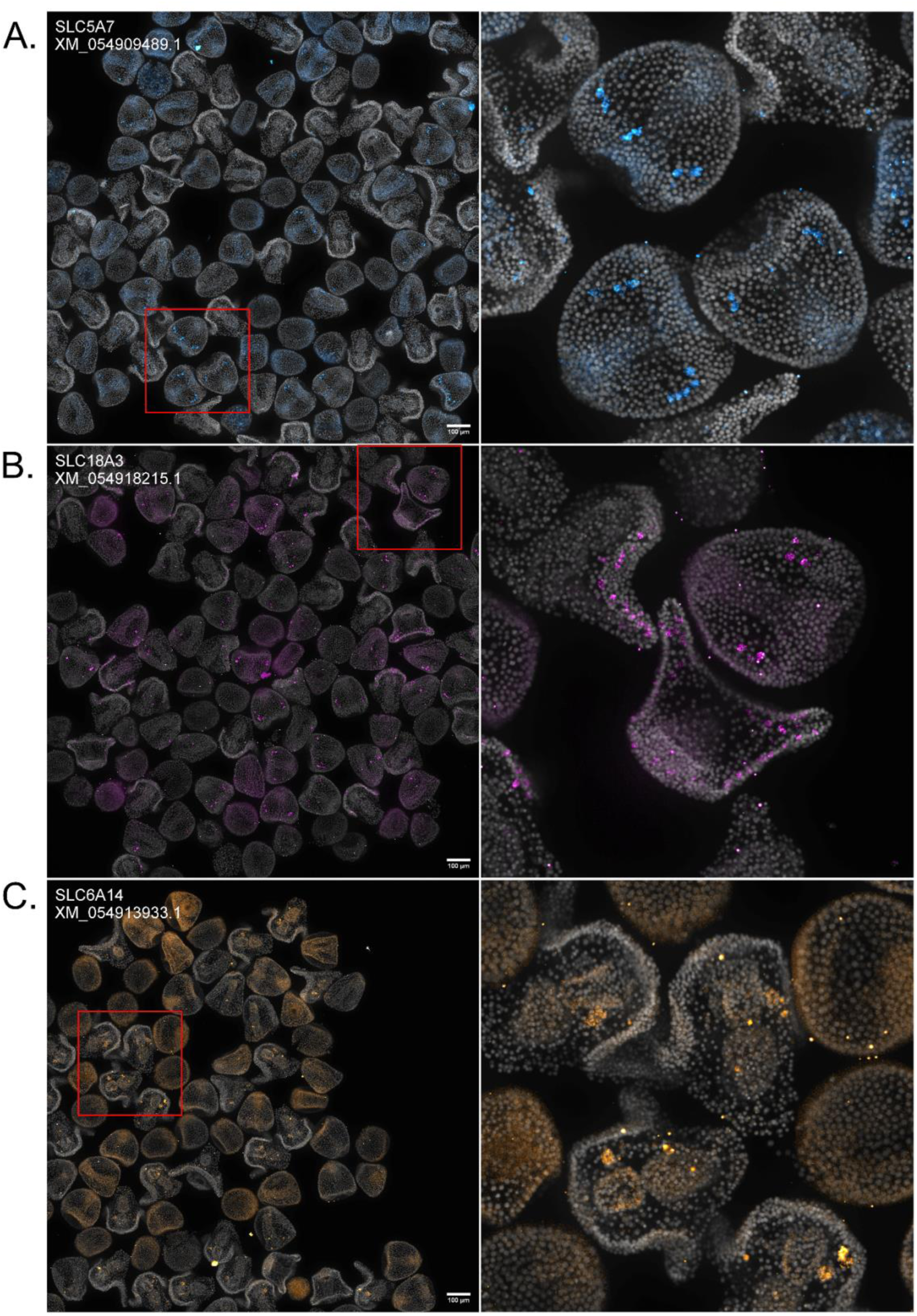
Novel SLC transporter expression patterns revealed by HT-HCR. All images contain a mix of 12, 24, and 36 hfp embryos and are automatically demultiplexed and stitched outputs resulting from an automated image processing workflow. Each image contains between 50-100 embryos and are annotated with the NCBI RefSeq ID for the mRNA sequence targeted by probe oligonucleotides. A) *SLC5A7*: Expression is either sparse or absent in 12 hfp embryos. At 24 hfp, expression is present in what appears to be neuronal precursor cells. At 36 hfp, expression is either sparse or absent. B) *SLC18A3*: Expression is either sparse or absent in 12 hfp embryos. At 24 hfp, expression is concentrated in in what appears to be neuronal precursor cells. Expression is concentrated in cells distributed throughout the ciliary band in 36 hfp embryos. C) *SLC6A14*: Expression is either sparse or absent in 12 hfp and 24 hfp embryos. At 36 hfp, expression is concentrated in one of the coelomic pouches.

We also localized other understudied membrane transporters critical to embryo survival and growth (**Figure 7**). One of these was *aquaporin-8 (Ishibashi et al., 1997; Koyama et al., 1997)*, a member of the aquaporin superfamily of membrane water channels (Agre and Kozono, 2003; King et al., 2004), which enables a rate of passage of water molecules across the cell membrane that is orders of magnitude higher than that of simple diffusion (Kozono et al., 2002; Law and Sansom, 2002). The expression of *aquaporin-8* is absent in 12 hpf embryos (**Figure 7A**). At 24 hpf expression is restricted to the premature hindgut and midgut and is excluded from the foregut (**Figure 7A**). The expression pattern is continued in 36 hpf embryos (**Figure 7A**). Finally, another important membrane protein that was localized was *short transient receptor potential channel 4* (*strp4*). *Strp4* is expressed in mesodermal cells (**Figure 7B**). At 12 hpf, expression is present in the NSM and at 24 and 36 hpf, expression is concentrated in mesodermal cells.

**Figure 7.**
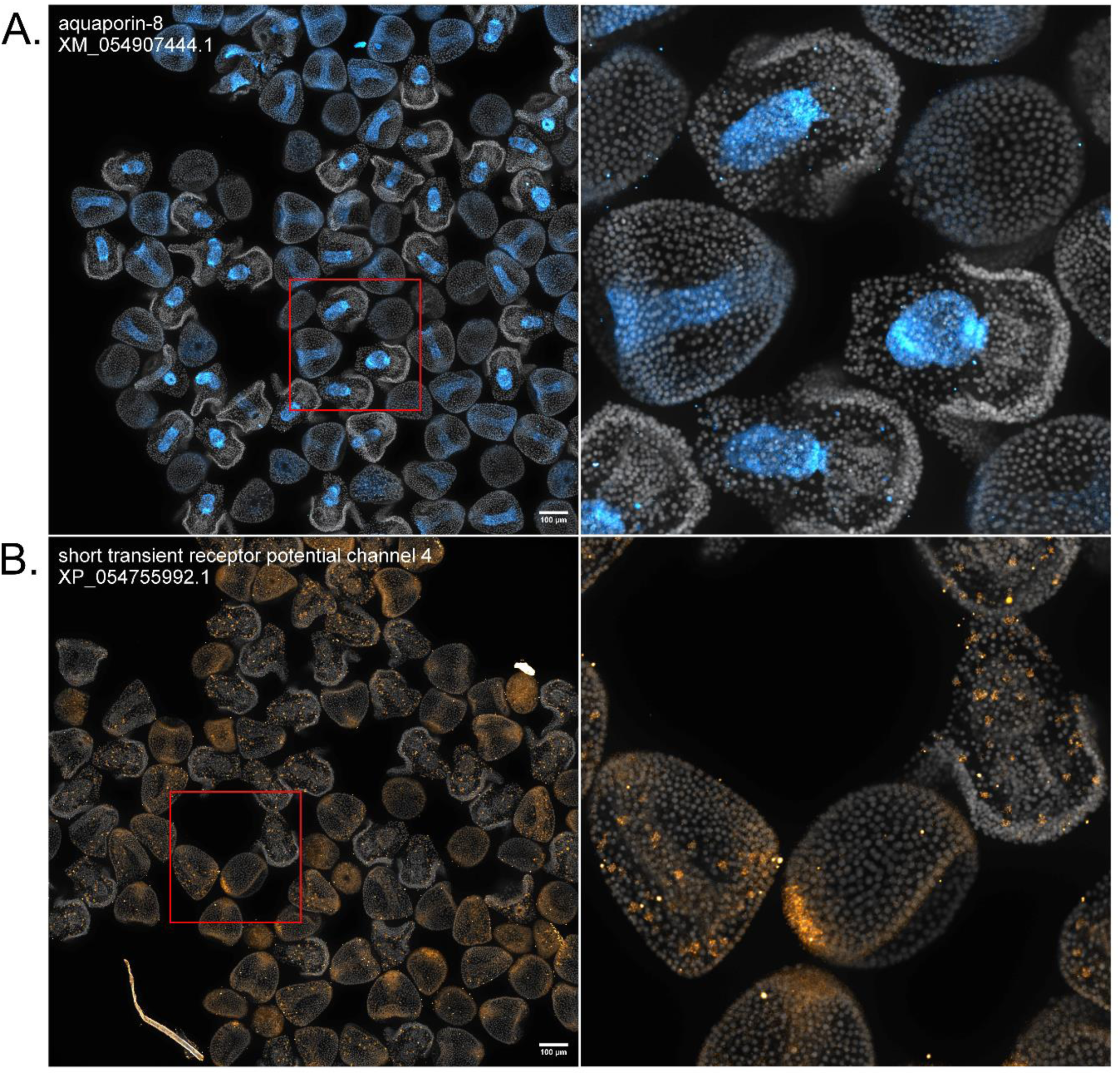
Aquaporin-8 and Strp-4 expression patterns revealed by HT-HCR. All images contain a mix of 12, 24, and 36 hfp embryos and are automatically demultiplexed and stitched outputs resulting from an automated image processing workflow. Each image contains between 50-100 embryos and are annotated with the NCBI RefSeq ID for the mRNA sequence targeted by probe oligonucleotides. A) *aquaporin-8*: Expression is either sparse or absent in 12 hfp embryos. At 24 hfp and 36 hfp, expression concentrated in the gut. B) *short transient receptor potential channel 4*: At 12 hfp, expression is concentrated in the NSM. At 24 hfp and 36 hfp, expression is concentrated in mesodermal cells.

## Discussion

For four decades, ISH has been a core methodology in the sea urchin community. While it is difficult to determine the exact number of genes localized, we estimate it to be <400 hundred genes. As demonstrated here, robotic liquid handling technologies and HCR have the potential to dramatically accelerate the pace of discovery using this animal model, and likely several others with similar biological features. Indeed, the localization of 101 genes in *L. pictus*, the first fully genetically enabled echinoderm (Jackson et al., 2024; Vyas et al., 2022), is itself an important foundational step for this emerging model.

### Features and limitations of HT-HCR

In general, HT-HCR is a high-throughput, automated, and miniaturized version of the established manual HCR protocol, and thus faces the same limitations related to reagent chemistry and interpretation of image data. As with manual approaches, not all genes targeted by HCR in our assay produced clearly interpretable images. In this case, from 192 targets, 101 provided clear expression patterns, while the other 91 lacked clear localizations (**Table S2**), potentially indicating weak or diffuse expression of the target gene. The images from these 91 ranged from appearing identical to negative controls to faint speckled signal. For example, there is a complete absence of signal for *MOB kinase 1A* (**Figure S4A, B**) consistent with negative controls, and faint, speckled signal for *titin* (**Figure S4C**), which may indicate weak ectodermal expression or simply be an unusual negative.

For genes with multiple paralogs, negative results could be a product of the specific paralogs or annotations chosen for probe design. Indeed, of these 91 probe sets, 15 were designed to target genes with multiple transcript variants. Consistent with this idea, for *mucin-2* and *mucin-17* two probe sets for these specific transcript variants showed clear expression patterns, while three (two for *mucin-2* and one for *mucin-17*) did not, and were thus considered part of the 91 negatives.

We speculated that manual HCRs might produce better quality images due to concerns involving the deterioration of reagent quality from sitting on the robotic deck at room temperature throughout the assay. However, as shown (**Figure 2**), we found no detectable difference in image quality between automatic runs and the manual runs. We were also concerned about whether the automated handling of embryos would harm the morphology of the samples. There was no clear indication of damage to or loss of embryos using automated pipetting, as the robotic protocol was designed to follow the same precautions taken during manual handling of embryo samples.

In this study, the HT-HCR protocol was used to process the commonly assayed developmental stages in the sea urchin research community. We expect that embryos in any developmental stage prior to 36 hpf will be viable for use with this method, as long as the fertilization envelope is removed for pre-hatching stages. However, the current HT-HCR protocol will require modification to handle larger embryos from later developmental stages (> 250um). This is because the protocol relies on the samples themselves being moved across different vessels throughout the assay, thereby enabling the use of highly miniaturized reaction volumes (and thus reductions of use of costly reagents, i.e. hairpins and amplifiers). A possible solution to this problem could be to use larger volumes, though this would increase the cost of the assay per sample and would still be limited by the maximum embryo size (∼0.5 mm) that can be moved by the handler. Automated protocols for larger samples such as late-stage larvae and juveniles, are unlikely to achieve the level of efficiency in reagent usage and imaging throughput presented here.

Finally, we also considered that many research projects in the echinoderm research community have been and may continue to be low throughput. Outside of large perturbation screens, the scale of HT-HCR may not be necessary for many applications. This said, our HT-HCR protocol was written for operation on a general-purpose laboratory liquid handler, and even small research groups may benefit from automated low or medium throughput processing of HCR samples because of the potential to reduce user-to-user variation.

### Implications and future directions

This study has several potential future implications for the use of ISH in sea urchins and related echinoderms. The first is that many more genes can now be described, including under-studied gene families. For example, the study of gene regulatory networks (GRNs) in urchins has been driven by extensive localization of the mRNA expression governed by transcription factors (TFs). However, many of the downstream effectors of these regulatory networks, such as the genes encoding structural or physiological proteins, remain under sampled. In addition, using HT-HCR or similar techniques, community level atlases of gene expression can quickly be acquired and linked to gene annotations (Telmer et al., 2024), as is the case for other animal models (Bradford et al., 2011; Fisher et al., 2023; Uhlen et al., 2010).

Perhaps more importantly because HT-HCR results in dramatic reduction in the time and cost per sample, it makes various types of perturbation analyses feasible. By nature of the readout, the results are directly relatable to the morphology of the embryo and thus, might provide insight into the mode of action of small molecule signals and teratogens. This could be particularly transformative for studying the effects of diverse small molecules on development using wild type, knockout (Tjeerdema et al., 2024) and transgenic urchins. These range from environmental chemicals (Anderson et al., 1994; Gambardella et al., 2021; Mijangos et al., 2020; Rendell-Bhatti et al., 2021; Sartori et al., 2023) to drugs (Gunaratne et al., 2018; Kim et al., 2022; Semenova et al., 2006; Tjeerdema et al., 2024) to endogenous molecules (Onjiko et al., 2015; Onjiko et al., 2016) - all of which have been speculated to act on development, but remain challenging to study at scale. To date, perturbation studies using ISH in urchins have necessarily focused on the expression of a limited subset of gene targets (Paganos et al., 2023; Rodríguez-Sastre et al., 2023). This approach opens the door to screening entire libraries of small molecule compounds against numerous genes, and thus could facilitate a new era of chemical biology using the sea urchin embryo.

Animal models have historically been selected because of their unique biological features (Hamdoun et al., 2023) that make them advantageous for certain avenues of investigation. In the case of the sea urchin, the biological features are extreme fecundity and developmental synchrony. This made it the model of choice for “biochemical scale” dissection of the molecular sequences of events occurring during development (Davidson et al., 2002; Evans et al., 1983). Yet as the field has transitioned from biochemical methods which utilize millions of embryos, to molecular and imaging-based methods which at most leveraged 100s of embryos, this advantage is diluted. This study demonstrates how the sea urchin biology may continue to be advantageous, in high-throughput, automated analysis of development.

## Acknowledgements

We thank Dr. Victor Vacquier for critical reading of the manuscript.

## Competing Interests

The authors declare no competing or financial interests.

## Funding

This work was supported by NIH ES035541, NSF 2414798, the Allen Discovery Center for Neurobiology in a Changing Environment and the Illumina Foundation.

## Data and resource availability

Scripts for generating probes and protocol file for HL_HTHCRv16 are available at https://github.com/yleesio/HL_HTHCR. Images for the 101 localizations can be viewed and downloaded at <u>Hamdoun Lab Image</u> <u>Repository</u>.

## Notes

https://github.com/ylee-sio/HL_HTHCR

https://drive.google.com/drive/folders/1gzPOIviqJyrCx1kx3u7cjfhPEX127gqp

### Competing Interest Statement

The authors have declared no competing interest.

