## Supplemental figures for "Automated, high-throughput in-situ hybridization of *Lytechinus pictus* embryos"

### Supplementary Material

**Figure S1.**

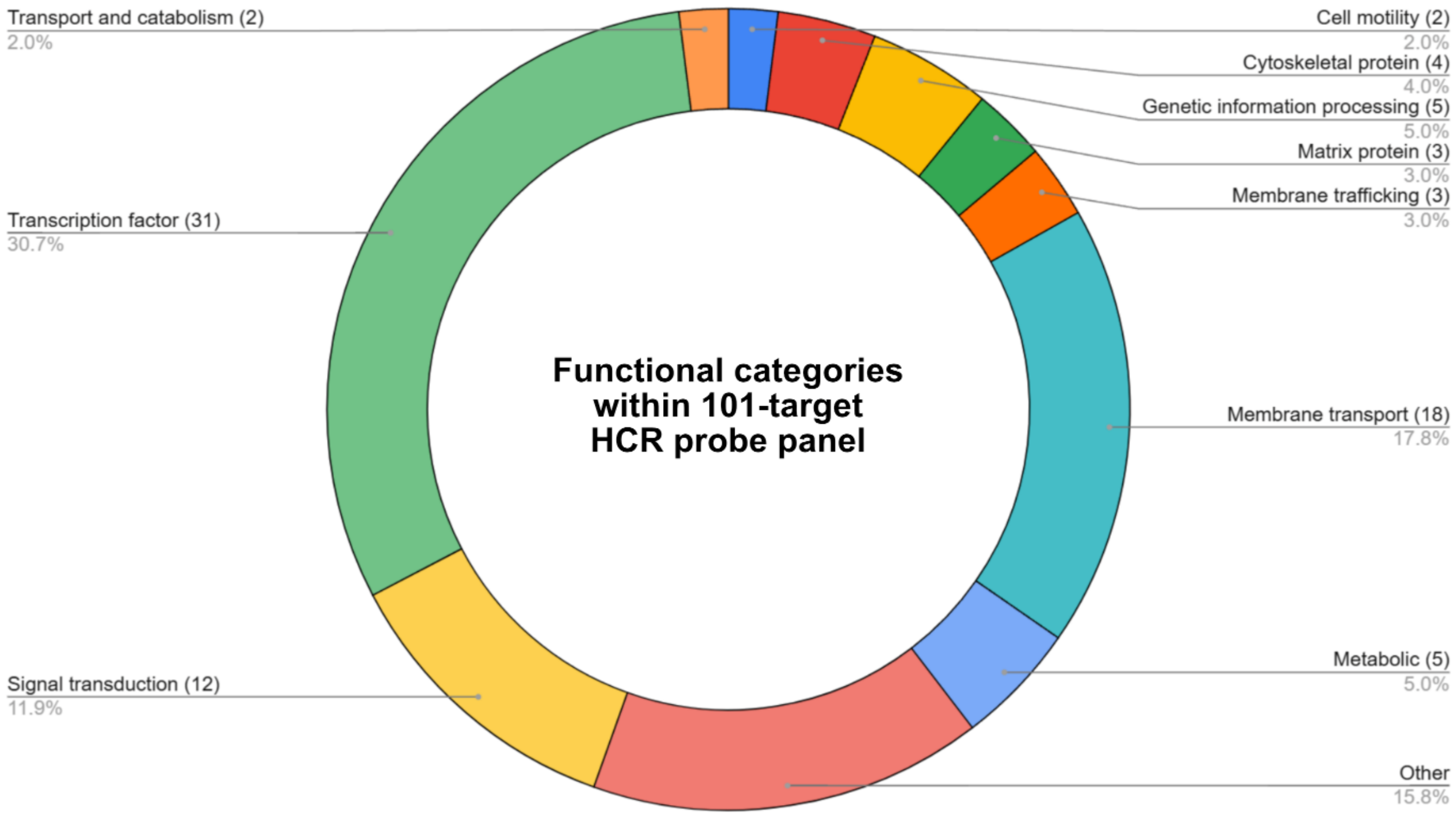
Transport and catabolism (2)
2.0%
Cell motility (2)
2.0%
Cytoskeletal protein (4)
4.0%
Genetic information processing (5)
5.0%
Matrix protein (3)
3.0%
Membrane trafficking (3)
3.0%
Transcription factor (31)
30.7%
Functional categories
within 101-target
HCR probe panel
Membrane transport (18)
17.8%
Metabolic (5)
5.0%
Signal transduction (12)
11.9%
Other
15.8%

**Figure S2.**

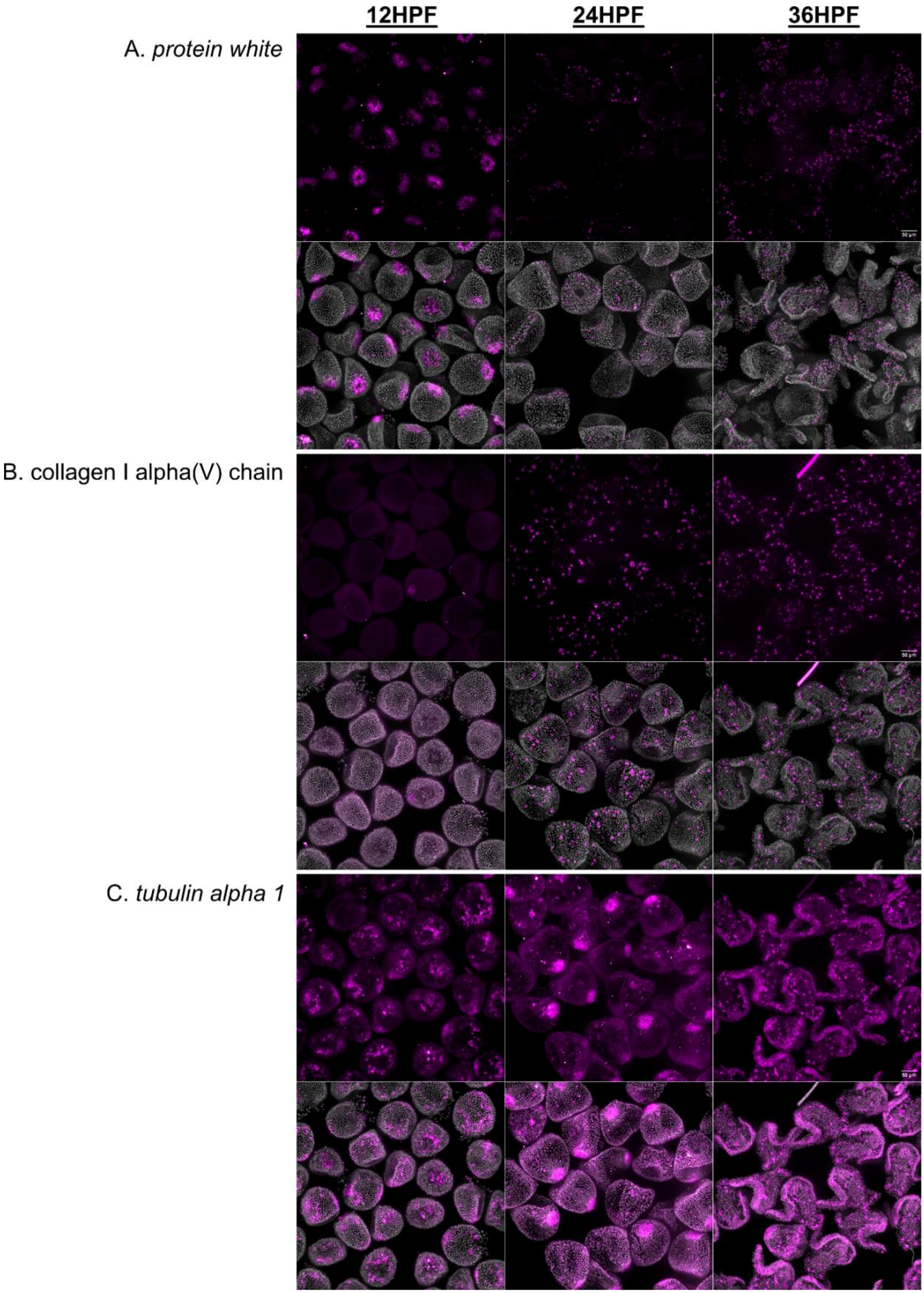
12HPF
24HPF
36HPF
A. *protein white*
B. collagen I alpha(V) chain
C. *tubulin alpha 1*

**Figure S3.**

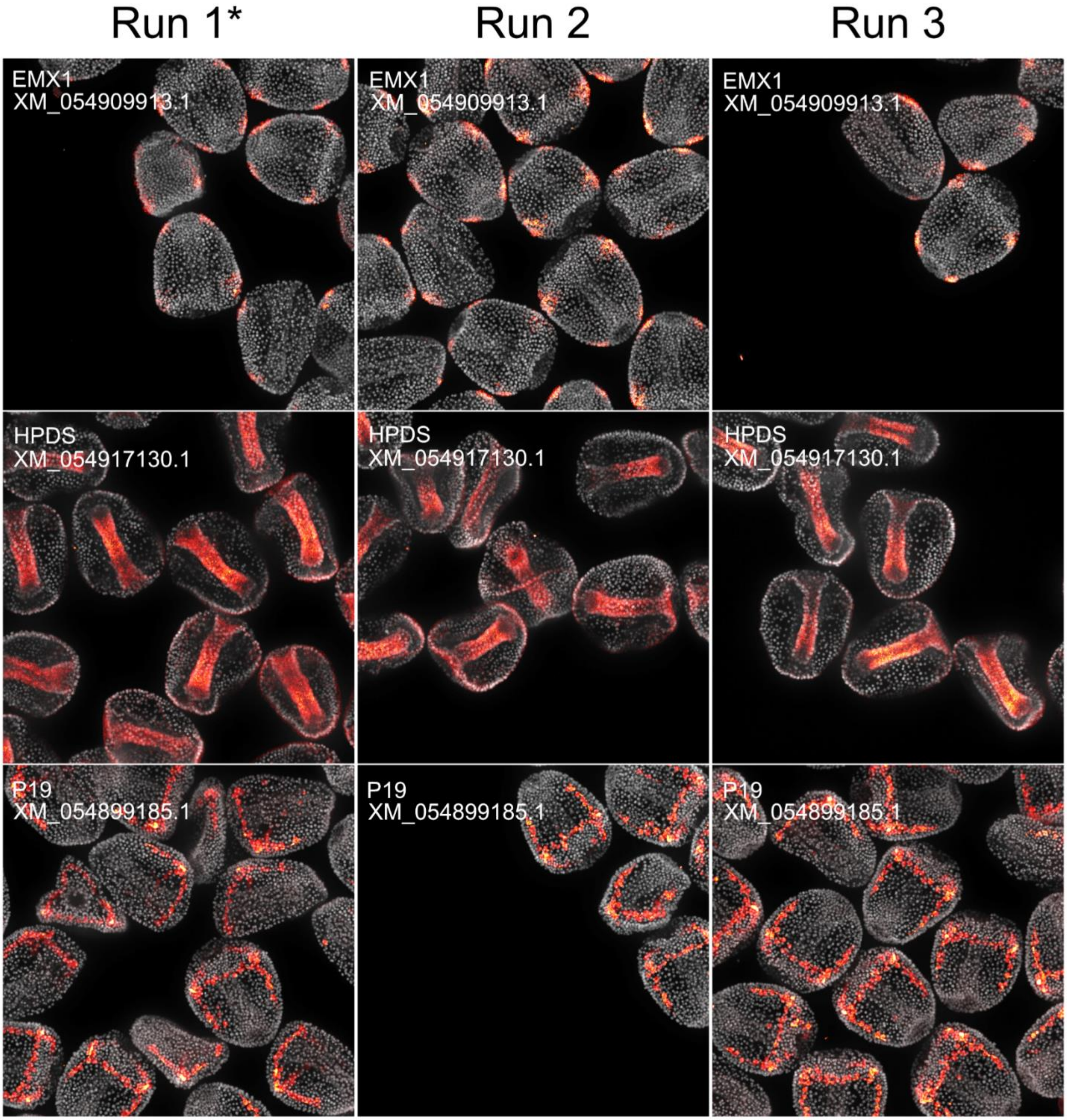

Run 1*
Run 2
Run 3
EMX1
XM_054909913.1
EMX1
XM_054909913.1
EMX1
XM_054909913.1
HPDS
XM_054917130.1
HPDS
XM_054917130.1
HPDS
XM_054917130.1
P19
XM_054899185.1
P19
XM_054899185.1
P19
XM_054899185.1

**Figure S4.**

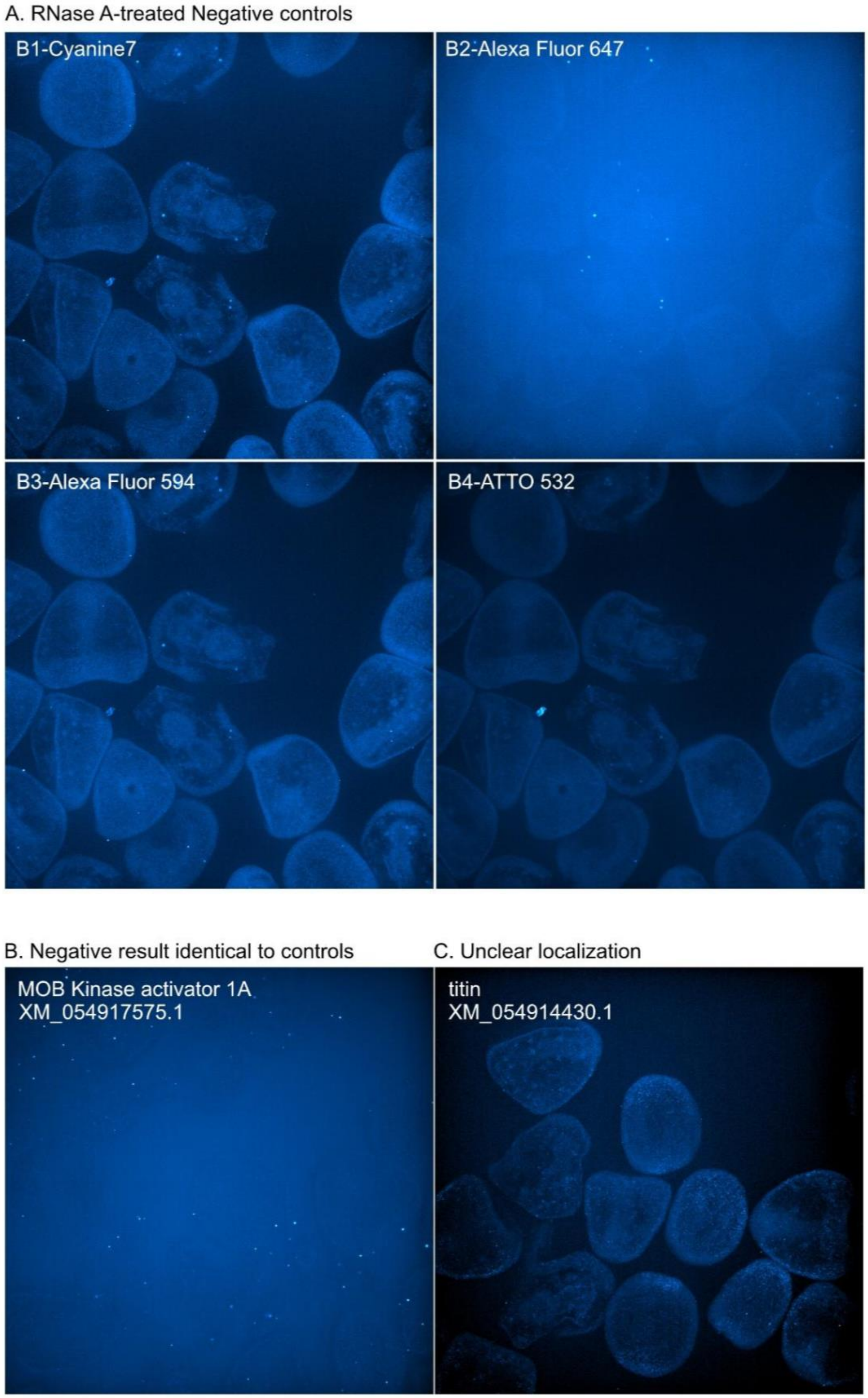
A. RNase A-treated Negative controls
B1-Cyanine7
B2-Alexa Fluor 647
B3-Alexa Fluor 594
B4-ATTO 532
B. Negative result identical to controls
MOB Kinase activator 1A
XM_054917575.1
C. Unclear localization
titin
XM_054914430.1

**Figure S1. Composition by gene functional category of the core HCR probe panel used in this study.**

**Figure S2. Automated sample processing for HCR produces clear localization patterns across different developmental stages.**

A) Localization of *protein white* (XM_054901325.1) in 12 hpf, 24 hpf, and 36 hpf embryos. In 12 hpf embryos, *protein white* is expressed in the NSM. In 24 and 36 hpf embryos, *protein white* is expressed in pigment cells. B) Localization of *collagen I alpha(V) chain* (XM_054913891.1) in 12 hpf, 24 hpf, and 36 hpf embryos. Expression is absent in 12 hpf embryos. In 24 and 36 hpf embryos, *collagen I alpha(V) chain* is expressed in mesodermal cells. C) Localization of *tubulin alpha 1* (XM_054897478.2) in 12 hpf, 24 hpf, and 36 hpf emrbyos. Expression of *tubulin alpha 1* is ubiquitous across all stages.

**Figure S3. Localization of genes are consistent across different runs of HT-HCR.**

Expression for *EMX1, HPDS,* and *P19* are consistent across three separate runs of HT-HCR. Asterisk (*) denotes images of genes which exist in another figure in this publication from the same run (Figure 2).

**Figure S4. RNAse A-treated negative control samples and genes which do not show clear localization.**

A) RNAse-treated negative control samples show no specific localization for the positive control probes (B1-*msp130*, B2-*FoxG*, B3-*calmodulin*, and B4-*chordin*) applied to the sample. B) Example of a gene (*MOB kinase activator 1A*; XM_054917575.1) whose signal appears identical to negative controls. C) Example of a gene (*titin*; XM_054914430.1) where signal is present but the localization is not clearly interpretable.

**Table S1. Genes which constitute 101 positive localizations in this study.**

| Accession | Gene Name | Gene Category |
|---:|---:|---:|
| XM_054913891.1 | collagen alpha-1(V) chain | cell_motility |
| XM_054897478.2 | tubulin alpha-1 chain | cell_motility |
| XM_054914077.1 | Actin, cytoskeletal 1A | cytoskeletal_protein |
| XM_054903584.1 | EMAP77 | cytoskeletal_protein |
| XM_054917843.1 | gelsolin-2 | cytoskeletal_protein |
| XM_054897076.1 | tubulin alpha chain-like | cytoskeletal_protein |
| XM_054915350.2 | 14-3-3 protein 2 | genetic_information_processing |
| XM_054901620.1 | APOBEC1 complementation factor | genetic_information_processing |
| XM_054908153.1 | ELAV3 | genetic_information_processing |
| XM_054902560.1 | elongation factor 1 gamma | genetic_information_processing |
| XM_054917449.1 | musashi1 | genetic_information_processing |
| XM_054914245.1 | ficolin-3 | matrix_protein |
| XM_054899185.1 | P19 | matrix_protein |
| XM_054919026.1 | SM34 | matrix_protein |
| XM_054906503.1 | ankyrin repeat domain-containing protein 55 | membrane trafficking |
| XM_064111875.1 | axotactin | membrane trafficking |
| XM_054918853.2 | synaptotagmin-1 | membrane trafficking |
| XM_054908194.1 | ABCA1 | membrane_transport |
| XM_054906925.1 | ABCA3 | membrane_transport |
| XM_054906809.1 | ABCA5 | membrane_transport |
| XM_054911547.1 | ABCB1 | membrane_transport |
| XM_064109812.1 | ABCB4 | membrane_transport |
| XM_054897890.1 | ABCB7 | membrane_transport |
| XM_054906725.1 | ABCC1 | membrane_transport |
| XM_054911957.1 | ABCG12 | membrane_transport |
| XM_054917858.1 | ABCG2 | membrane_transport |
| XM_054907444.1 | aquaporin-8 | membrane_transport |
| XM_054901325.1 | protein_white | membrane_transport |
| XM_064095033.1 | short transient receptor potential channel 4 | membrane_transport |
| XM_054900099.1 | SLC13A5 | membrane_transport |
| XM_054910749.1 | SLC18A2 | membrane_transport |

| | | |
|---|---|---|
| XM_054918215.1 | SLC18A3 | membrane_transport |
| XM_054913204.1 | SLC2A1 | membrane_transport |
| XM_054909489.1 | SLC5A7 | membrane_transport |
| XM_054913933.1 | SLC6A14 | membrane_transport |
| XM_054900733.1 | betaine-homocysteine S-methyltransferase 1 | metabolic |
| XM_054918411.1 | D-dopachrome decarboxylase | metabolic |
| XM_054916386.1 | eyes absent | metabolic |
| XM_054914495.1 | hematopoietic prostaglandin D synthase (HPDS) | metabolic |
| XM_054915559.1 | quinone oxidoreductase | metabolic |
| XM_054900056.1 | beta-catenin | other |
| XM_054915079.1 | collagen alpha-3(IV) chain/COLP3alpha | other |
| XM_054895445.2 | COLP1alpha | other |
| XM_054892573.1 | cyclophilin-1 | other |
| XM_054919060.1 | Endo16 | other |
| XM_064102407.1 | fatty acid-binding protein type 3 | other |
| XM_054900255.1 | matrix metalloproteinase-2 | other |
| XM_054896825.1 | metallothionein-A | other |
| XM_054912810.1 | msp130 | other |
| XM_054893990.1 | mucin-17 | other |
| XM_054899132.1 | mucin-2 | other |
| XM_054901731.1 | protein PB18E9.04c | other |
| XM_054911221.1 | protein rolling stone | other |
| XM_054899164.1 | protocadherin Fat 1-like | other |
| XM_054902877.1 | Spec3 | other |
| XM_054905379.1 | troponin I | other |
| XM_054917286.1 | bmp2 | signal_transduction |
| XM_054900839.1 | calmodulin | signal_transduction |
| XM_054906303.1 | CD151 antigen | signal_transduction |
| XM_054912384.1 | chordin | signal_transduction |
| XM_054911692.1 | dickkopf WNT signaling pathway inhibitor 3-like | signal_transduction |
| XM_054897154.1 | frizzled-4 | signal_transduction |
| XM_054897832.1 | frizzled-5 | signal_transduction |
| XM_054902120.1 | left-right determination factor 2 | signal_transduction |

| | | |
|---|---|---|
| XM_064102363.1 | neuroblast differentiation-associated protein AHNAK-like | signal_transduction |
| XM_054908715.1 | patched homolog 1-like | signal_transduction |
| XM_054902965.1 | univin | signal_transduction |
| XM_054917943.1 | wnt8 | signal_transduction |
| XM_054897283.1 | blimp1a | transcription_factor |
| XM_054899987.1 | dri | transcription_factor |
| XM_054904347.1 | erg | transcription_factor |
| XM_054905949.1 | ese | transcription_factor |
| XM_054910758.1 | transcription factor coe | transcription_factor |
| XM_054903096.1 | FoxA | transcription_factor |
| XM_054892407.1 | FoxC | transcription_factor |
| XM_054903002.1 | FoxG | transcription_factor |
| XM_054906495.1 | FoxQ2 | transcription_factor |
| XM_054916700.2 | gcm | transcription_factor |
| XM_054895438.1 | Hbox7 | transcription_factor |
| XM_054909913.1 | homeobox protein EMX1 | transcription_factor |
| XM_054900719.1 | homeobrain | transcription_factor |
| XM_054892815.1 | odd-skipped-related 1 | transcription_factor |
| XM_054901727.1 | onecut2 | transcription_factor |
| XM_054917528.1 | pax2/5/8 | transcription_factor |
| XM_054896981.1 | POU domain, class 3, transcription factor | transcription_factor |
| XM_054902058.1 | prospero homeobox protein 1 | transcription_factor |
| XM_054912035.1 | ptfa1 | transcription_factor |
| XM_054913367.1 | scratch | transcription_factor |
| XM_054895154.1 | fezf2 | transcription_factor |
| XM_054902743.1 | six1 | transcription_factor |
| XM_054902744.1 | six3/6 | transcription_factor |
| XM_054914671.1 | Sox-3-B | transcription_factor |
| XM_054898009.1 | T-box brain transcription factor 1 | transcription_factor |
| XM_054897143.1 | T-box transcription factor T (brachyury) | transcription_factor |
| XM_054902833.1 | thyroid transcription factor 1 | transcription_factor |
| XM_054898855.1 | thyrotroph embryonic factor-like | transcription_factor |
| XM_054906577.1 | transcription factor SOX-9 | transcription_factor |

| | | |
|---|---|---|
| XM_054895436.1 | XHOX-3 | transcription_factor |
| XM_054897132.1 | zinc finger C4H2 domain contaning protein | transcription_factor |
| XM_054896288.1 | caveolin-3 | transport_and_catabolism |
| XM_054895022.1 | ManR | transport_and_catabolism |

**Table S2. Genes which failed to produce clearly interpretable localization.**

| Accession | Gene Name |
|---|---|
| XM_054913519.1 | abca2 |
| XM_054916213.1 | abca4 |
| XM_054902369.1 | abcb10 |
| XM_054911894.1 | abcb6 |
| XM_054893952.1 | abcb9 |
| XM_054893102.1 | abcc10a |
| XM_054902028.1 | abcc10b |
| XM_054896220.1 | ABCC9 |
| XM_054903689.1 | adipocyte plasma membrane-associated protein |
| XM_054913502.1 | allograft inflammatory factor 1-like |
| XM_054900595.1 | ameboid myosin I |
| XM_054898670.1 | aryl_hydricarbon_receptor_nuclear_translocator_homolog |
| XM_054906884.1 | brain-specific angiogenesis inhibitor 1-associated protein 2 |
| XM_054915270.1 | BRCA2-interacting_transcriptional_repressor_EMSY-like |
| XM_054893801.1 | caspase_6_like |
| XM_054919029.1 | catalase-like |
| XM_054905955.1 | CD9 antigen |
| XM_054892002.1 | coronin-1B |
| XM_054904859.1 | cubilin-like |
| XM_054906733.1 | DNA ligase 1 |
| XM_054910483.1 | drebrin-like protein A |
| XM_054893174.1 | hnf1a |
| XM_041598698.1 | hnf4 |
| XM_054901951.1 | homeobox protein homothorax-like |
| XM_054907821.1 | homeobox protein Unc−4 |
| XM_054894536.1 | hox1 |
| XM_054896340.1 | hox9 |
| XM_054906011.1 | hsp70-binding_protein_1_like |
| XM_054919221.1 | hsp90_co_chaperone_cdc_37_like |
| XM_054914601.1 | hyalin-like |
| XM_054899136.1 | Kibra-like |

| | |
|---|---|
| XM_054907292.1 | lethal(2) giant larvae protein homolog 1-like |
| XM_054897344.1 | macoilin |
| XM_054908910.1 | macrophage migration inhibitory factor |
| XM_054897977.1 | metal_regulatory_transcription_factor_1_like |
| XM_054913983.1 | metal_response_element_binding_transcription_factor_2_like |
| XM_054906133.1 | microtubule-associated_protein_futsch-like |
| XM_054917575.1 | MOB kinase activator 1A-like |
| XM_054906288.1 | MOB kinase activator 2-like |
| XM_054897489.1 | mtf1 |
| XM_054893604.1 | mucin-5ac |
| XM_054899059.1 | mucin-12 |
| XM_054909363.1 | mucin-19 |
| XM_054905261.1 | mucinl |
| XM_054919698.1 | neuronal acetylcholine receptor subunit alpha-10 |
| XM_054912738.1 | neuronal acetylcholine receptor subunit alpha-2 |
| XM_054915363.1 | neurotrypsin |
| XM_054899223.1 | octopamine receptor |
| XM_054901330.1 | plastin-3 |
| XM_054902250.1 | protein salvador homolog 1-like |
| XM_054904308.1 | remodeling and spacing factor 1 |
| XM_054899962.1 | retina and anterior neural fold homeobox protein |
| XM_054904870.1 | roundabout homolog 1 |
| XM_054899015.1 | secretory carrier-associated membrane protein 1 |
| XM_054916437.1 | Serine/threonine-protein kinase 3-like (hippo) |
| XM_054899570.1 | Serine/threonine-protein kinase 4-like (hippo) |
| XM_054896280.1 | serine/threonine-protein kinase A-Raf-like |
| XM_054897360.1 | serine/threonine-protein kinase LATS2-like |
| XM_054896684.1 | slc22a4 |
| XM_054913357.1 | slc31a1 |
| XM_054898789.1 | smad1/5/8 |
| XM_054906962.1 | SOCS2 |
| XM_054903022.1 | spastin |
| XM_054900002.1 | sprouty homolog 3-like |

| | |
|---|---|
| XM_054907292.1 | sushi |
| XM_054919067.1 | synaptotagmin-7 |
| XM_054895192.1 | TAZ |
| XM_054918745.1 | TEF1-like |
| XM_054893323.1 | tektin1 |
| XM_054914430.1 | titin |
| XM_054904575.1 | tyrosine-protein kinase JAK2 |
| XM_054919211.1 | tyrosine-protein_phosphatase_10D-like |
| XM_054894772.1 | wnt16 |
| XM_054915720.1 | YAP1-A |
| XM_054914833.1 | YAP1-B |
| XM_054909721.1 | z60/egr |
| XM_054913988.1 | abcc4_X1_2* |
| XM_054915178.1 | abcc4_X1* |
| XM_054915179.1 | abcc4_X2* |
| XM_054915180.1 | abcc4_X3* |
| XM_054915181.1 | abcc4_X4* |
| XM_054900846.1 | caspase_3_like* |
| XM_054897466.1 | caspase_3_like* |
| XM_054896836.1 | cytochrome_P450_1A1_like* |
| XM_054904951.1 | cytochrome_P450_1A1_like* |
| XM_054906659.1 | cytochrome_P450_1A1_like* |
| XM_054893754.1 | mucin-17_like* |
| XM_054905346.1 | mucin-17_like* |
| XM_054905746.1 | mucin-2_like* |
| XM_054892398.1 | mucin-2_like* |
| XM_054895498.1 | mucin-2_like* |
| | * denotes transcript isoform for the same gene |
